# Systemic, RPE-directed AAV-*Tyrosinase* therapy restores ocular pigmentation in an OCA1 mouse model

**DOI:** 10.64898/2026.07.01.735814

**Authors:** Alessandra M. Larimer-Picciani, Lucas B. Jacob, Kacey J. Sullinger, Wyatt G. Kriebel, José-Alain Sahel, Leah C. Byrne

## Abstract

Oculocutaneous albinism type 1 (OCA1) is a pigmentation disorder caused by biallelic *tyrosinase* (*TYR*) mutations, an essential enzyme for melanin synthesis. TYR inactivity results in loss of hair, skin, and eye pigment, which is detrimental for ocular function. Hypopigmentation of iris, retinal pigment epithelium (RPE), and choroid results in severe photosensitivity and low visual acuity. There are currently no FDA-approved pigment restoring therapies for OCA1, making therapeutic development an unmet clinical need. To address this gap, we have advanced an adeno-associated viral (AAV)-mediated *Tyr* replacement approach for OCA1 ocular pigment restoration. We evaluated the optimal viral delivery strategy and vector cell-type specificity for iris, RPE, and choroid pigmentation in an OCA1 mouse model, testing intraocular and systemic viral delivery methods in conjunction with viral constructs of varying RPE-specificity. Early, systemic delivery of an RPE-directed AAV-*Tyr* construct, AAV9.2yf-VMD2-*Tyr*, achieved widespread ocular pigment rescue with minimal off-target expression in non-ocular tissues. Animals treated with AAV9.2yf-VMD2-*Tyr* demonstrated reduced photophobic behavior compared to untreated controls, indicating that ocular pigmentation restores a debilitating functional consequence of OCA1. Our findings establish a foundation for clinical translation of an AAV-*TYR* therapy aimed at improving light sensitivity, glare, and low vision through pigment restoration in patients with OCA1.

## INTRODUCTION

Oculocutaneous albinism (OCA) is a genetic disorder characterized by reduced pigment production in the hair, skin, and eyes. OCA is caused by a heterogenous group of autosomal recessive mutations that disrupt melanin synthesis or storage and affects approximately 1 in 17,000 individuals worldwide^1^. Oculocutaneous albinism type 1 (OCA1) is the most common OCA subtype (1:36,000)^2,3^ and is caused by biallelic mutations in *tyrosinase* (*TYR*), a melanogenic enzyme responsible for the initial and rate limiting steps in melanin synthesis. While the clinical severity of OCA1 can vary with residual TYR function^4^, the complete loss of TYR activity results in the most profound hypopigmentation phenotype among all eight non-syndromic OCA subtypes^5^.

Loss of pigment and TYR function is particularly detrimental to ocular function. *TYR* is expressed in multiple ocular tissues, including the iris, retinal pigment epithelium (RPE), and choroid^6^. Pigment in these tissues absorbs excess ultra-violet light, protecting photoreceptors from light-induced damage, and reduces light scatter, thereby reducing visual glare^7^. TYR and pigment is particularly important within the RPE^8^. In addition to absorbing stray light, melanin within the RPE acts as a free radical scavenger, protecting interdigitated photoreceptors from oxidative stress^9^, and facilitates photoreceptor outer segment recycling, delaying age-related photoreceptor degeneration^7,10^.

In addition to its photoprotective roles, *TYR* expression and RPE pigmentation are essential for retinal development^11,12^. Through mechanisms that are incompletely understood, *TYR* deficiency leads to retinal ganglion cell (RGC) misrouting and foveal hypoplasia^13,14^. Together, ocular hypopigmentation and retinal developmental anomalies result in photophobia, glare, abnormally low visual acuity, and impaired stereoscopic vision; these vision-limiting symptoms are often further compounded by nystagmus, strabismus, and amblyopia in patients with OCA1^15^.

Current OCA1 clinical management focuses on sun protection and optimizing residual visual function^16^. Patients are at an increased risk for basal and squamous cell carcinomas^17^, making cutaneous sun protection essential. Tinted contact lenses or dark spectacles may be used for ocular light protection; however, these measures offer only partial relief from photophobia/glare and may worsen already reduced contrast sensitivity^16^. Low visual acuity is managed through corrective lenses, low vision aids, and rehabilitation, but remains limited by retinal maldevelopment and persistent photosensitivity. There are currently no FDA-approved therapies to increase ocular pigment and ameliorate the visual effects of TYR-negative OCA1^18^, making therapeutic development an urgent unmet clinical need.

OCA1 is an ideal disease candidate for gene replacement therapy. One functional copy of *TYR* is sufficient to restore ocular pigmentation and prevent the most debilitating symptoms associated with OCA1, including photosensitivity and low visual function^19^, and the eye is easily accessible for gene therapy delivery^20^. Adeno-associated virus (AAV) is the preferred vector for ocular gene replacement. AAVs have demonstrated excellent efficacy and safety in human ocular gene therapy trials, as exemplified by Luxturna, an FDA-approved AAV-based therapy for RPE65-deficient Leber’s congenital amaurosis^21,22^.

Defining the AAV-*TYR* delivery strategy that maximizes ocular pigment restoration is a necessary step toward the development of a clinically viable therapy. Proof-of-concept studies by Gargiulo et al. demonstrated that adult AAV-*Tyr* therapy can partially restore RPE and choroid pigmentation in an OCA1 mouse model^23^. While this work established the feasibility of AAV-*Tyr* ocular pigment rescue, it did not address key variables that influence therapeutic success, including the optimal method of delivery or viral cell-type specificity. Given the roles of ocular pigmentation in minimizing intraocular light scatter, reducing visual glare, and supporting visual function, we hypothesize that the magnitude of pigment restoration in the RPE, choroid, and iris will correlate with greater functional benefit. Thus, identifying the delivery method and transgene expression specificity that confer the most robust pigment restoration in these tissues is essential in advancing AAV-*TYR* clinical translation.

In this study, we systematically evaluated AAV-*Tyr* delivery strategies for restoring ocular pigmentation in a validated OCA1mouse model^24^. We tested three AAV delivery methods, including systemic (intravenous) and intraocular (subretinal and intravitreal) modes of viral delivery, and compared a ubiquitous versus an RPE-promoter to evaluate the effect of transgene expression specificity on pigment production. Our findings demonstrate that early, systemic delivery of an RPE-directed *Tyr* construct resulted in the most effective and targeted pigmentation rescue, with minimal off-target expression in non-ocular tissues. Treated animals exhibited reduced photophobic behavior compared to untreated controls, AAV-mediated ocular pigment restoration ameliorates a debilitating consequence of OCA1.

## RESULTS

To evaluate how the method of recombinant AAV (rAAV) delivery affects ocular pigment restoration, we administered the ubiquitously expressed rAAV-CAG-*Tyr* construct using three delivery routes: intravenous injection via the facial vein and two common intraocular techniques, subretinal and intravitreal injection (Figure 1A, Figure S1). The AAV9.2yf capsid was used for intravenous and subretinal delivery due to its enhanced ability to traverse vascular endothelium and transduce the RPE^25^. An engineered AAV2-based capsid developed from our state-of-the-art viral engineering platform^26^ was used to bypass the inner limiting membrane (ILM) for intravitreal delivery.

**Figure 1.**
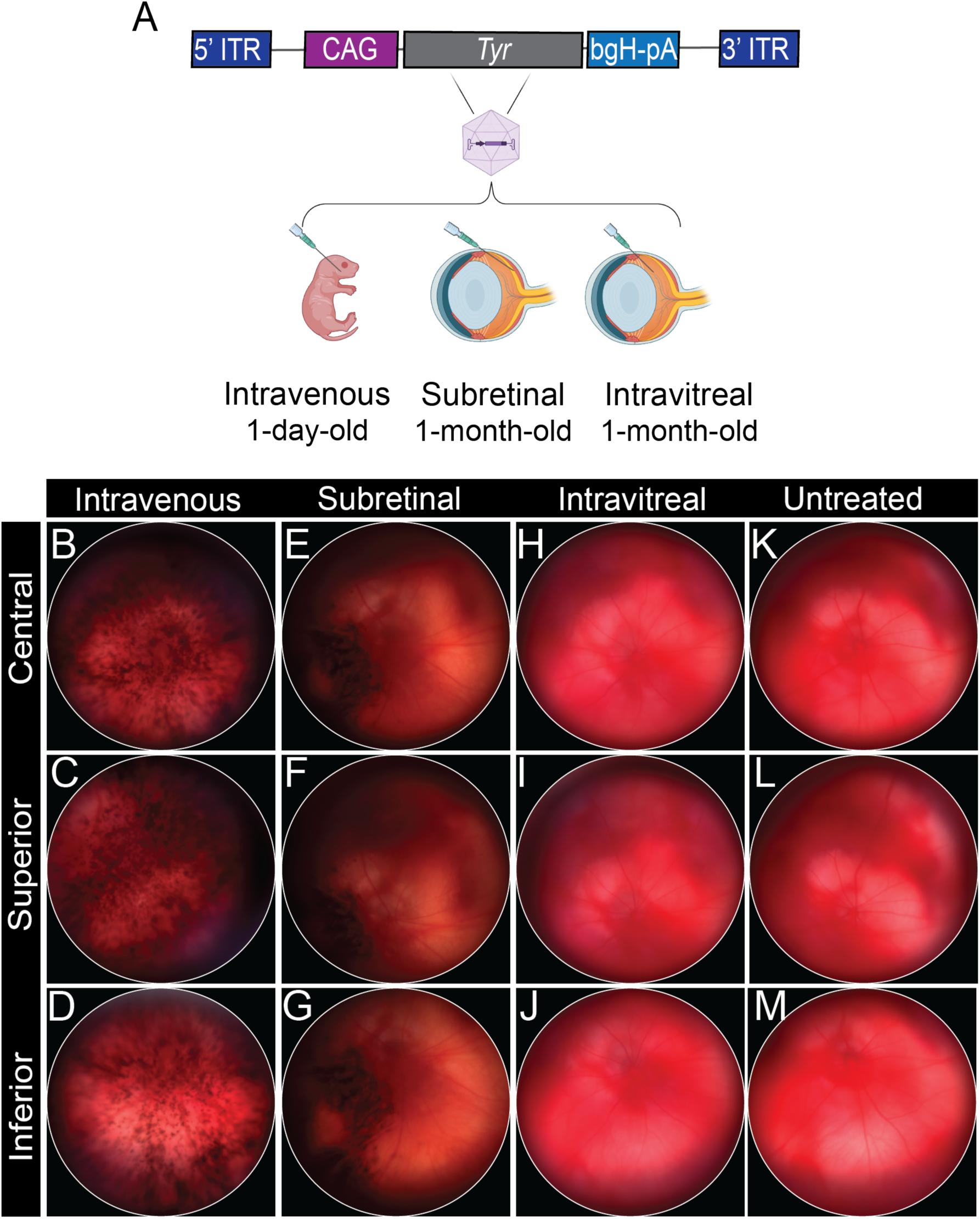
Intravenous and subretinal AAV9.2yf-CAG-*Tyr* restored OCA1 posterior segment melanization with delivery-dependent pigmentation patterns. (A) Schematic overview of the self-complementary (sc) ubiquitous AAV vector transgene (scAAV-CAG-*Tyr*) used for intravenous, subretinal, and intravitreal injection. 50 uL of 2.5e12 vector genome (vg)/mL AAV9.2yf-scCAG-*Tyr* was delivered via the facial vein in P1 mice. Subretinal and intravitreal injection of 1.5 uL 2.5e12 vg/mL AAV9.2yf-scCAG-*Tyr* was performed in one-month-old mice. (B-J) *In vivo* fundus images acquired from mice one-month post viral delivery. (B-D) P1 intravenous AAV9.2yf-scCAG-*Tyr* resulted in widespread, mottled pigment production throughout the posterior segment, underlying the central (B), superior (C) and inferior (D) retina. (E-G) Subretinal AAV9.2yf-scCAG-*Tyr* restored pigment around the site of injection, underlying the inferior (G) retina. Intravitreal delivery did not result in visible superior (H), central (I), or inferior (J) posterior segment pigmentation. Central (K), superior (L), and inferior (M) retinal images of untreated B6(Cg)-Tyr^c-2J^/J mice; posterior segment pigmentation is absent.

Intravenous injections were performed at post-natal day 1, prior to the formation of the blood-retina barrier^27^, while intraocular injections were administered in 1-month-old OCA1 mice. *In vivo* fundus imaging performed one-month post-viral delivery revealed that intravenous delivery of AAV9.2yf-CAG-*Tyr* resulted in widespread, mottled pigmentation rescue across the posterior segment (Figure 1B-D). In contrast, subretinal delivery led to localized pigmentation surrounding the injection site (Figure 1E-G). No posterior segment pigmentation was observed following intravitreal delivery at the dose tested (3.8e9 vg) (Figure 1H-J). Histological analysis confirmed melanin localization within the iris, RPE, and choroid. Immunohistochemical staining demonstrated melanin-associated Tyr localization (Figure S2).

To determine whether RPE-directed expression could achieve comparable posterior segment pigment restoration, we tested the *VMD2* promoter^28^ (Figure 2A, Figure S1). As with the ubiquitous construct, we evaluated pigmentation following intravenous, subretinal, and intravitreal delivery using the same capsids described above. Intravenous administration of AAV9.2yf-VMD2-*Tyr* at P1 resulted in widespread, mottled pigmentation across all quadrants of the posterior segment (Figure 2B-D), similar to the pigment restoration pattern achieved with the CAG promoter, but with modestly greater posterior segment pigment coverage compared to the ubiquitous vector, qualitatively. The subretinal approach again yielded localized pigmentation restricted to the site of neurosensory detachment induced by the injection (Figure 2E-G). No posterior segment pigmentation was observed following intravitreal delivery (Figure 2H-J). Histological analysis revealed pigment restoration in the iris, RPE, and choroid. Immunohistochemical staining demonstrated melanin-associated Tyr localization (Figure S2).

**Figure 2.**
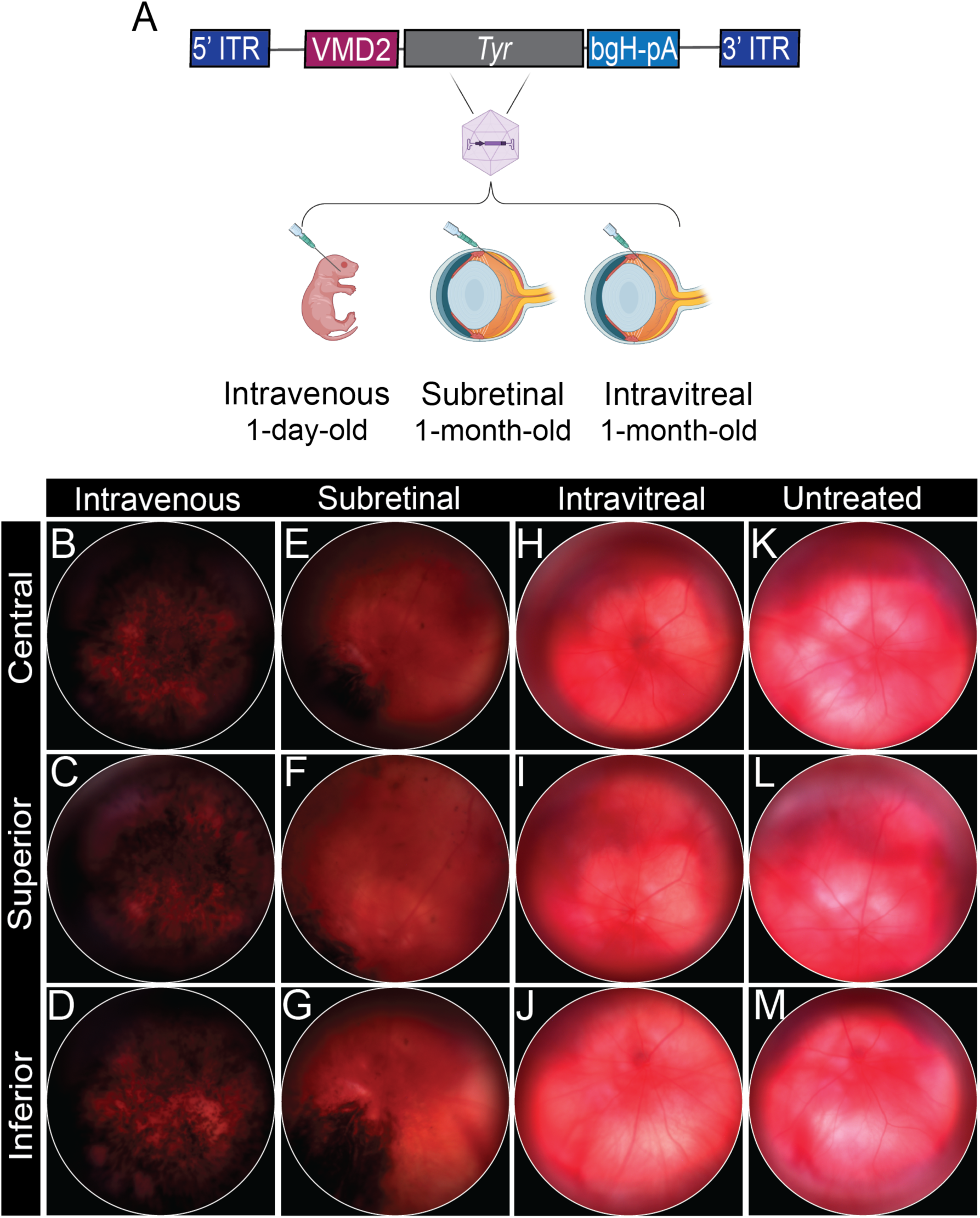
The RPE-directed vector AAV9.2yf-VMD2-*Tyr* restored OCA1 posterior segment pigmentation with delivery-dependent melanin deposition patterns. (**A**) Schematic overview of the RPE-selective AAV vector transgene (scAAV-VMD2-*Tyr*) used for intravenous, subretinal, and intravitreal injection. 50 uL of 2.5e12 vector genome (vg)/mL AAV9.2yf-scVMD2-*Tyr* was delivered via the facial vein in P1 mice. Subretinal and intravitreal injection of 1.5 uL 2.5e12 vg/mL AAV9.2yf-scVMD2-*Tyr* was performed in one-month-old mice. (**B-J**) *In vivo* fundus images acquired from mice one-month post viral delivery. (**B-D**) P1 intravenous AAV9.2yf-scVMD2-*Tyr* resulted in widespread, mottled pigment production throughout the posterior segment, underlying the central (**B**), superior (**C**) and inferior (**D**) retina. (**E-G**) Subretinal AAV9.2yf-scVMD2-*Tyr* restored pigment around the site of injection, concentrated over the inferior (**G**) retina and resulting in peripheral mottling overlying the superior (**E**) retina. Intravitreal delivery did not result in visible superior (**H**), central (**I**), or inferior (**J**) posterior segment pigmentation. Central (**K**), superior (**L**), and inferior (**M**) retinal images of untreated B6(Cg)-Tyr^c-2J^/J mice; posterior segment pigmentation was absent.

To more comprehensively assess anterior and posterior segment pigmentation across delivery methods, we performed whole-eye clearing using eumelanin-sparing methods^29^. This approach revealed the same posterior segment pigmentation patterns visualized on *in vivo* imaging and enabled iris pigment visualization.

Intravenous AAV9.2yf-CAG-*Tyr* and AAV9.2yf-VMD2-*Tyr* both restored pigment throughout the posterior segment, as well as partial iris pigmentation (Figure 3A-F). Again, the VMD2 promoter appeared to result in modestly greater pigment production compared to the ubiquitous construct, particularly following intravenous injection, qualitatively. In contrast, subretinal delivery of both constructs resulted in pigmentation concentrated at the injection site, with sparse distal pigmentation and minimal iris involvement (Figure 3G-L). In corroboration with *in vivo* imaging, intravitreal delivery did not restore posterior segment pigmentation in whole eyes. However, dense iris pigmentation was observed following CAG-*Tyr* treatment.

**Figure 3.**
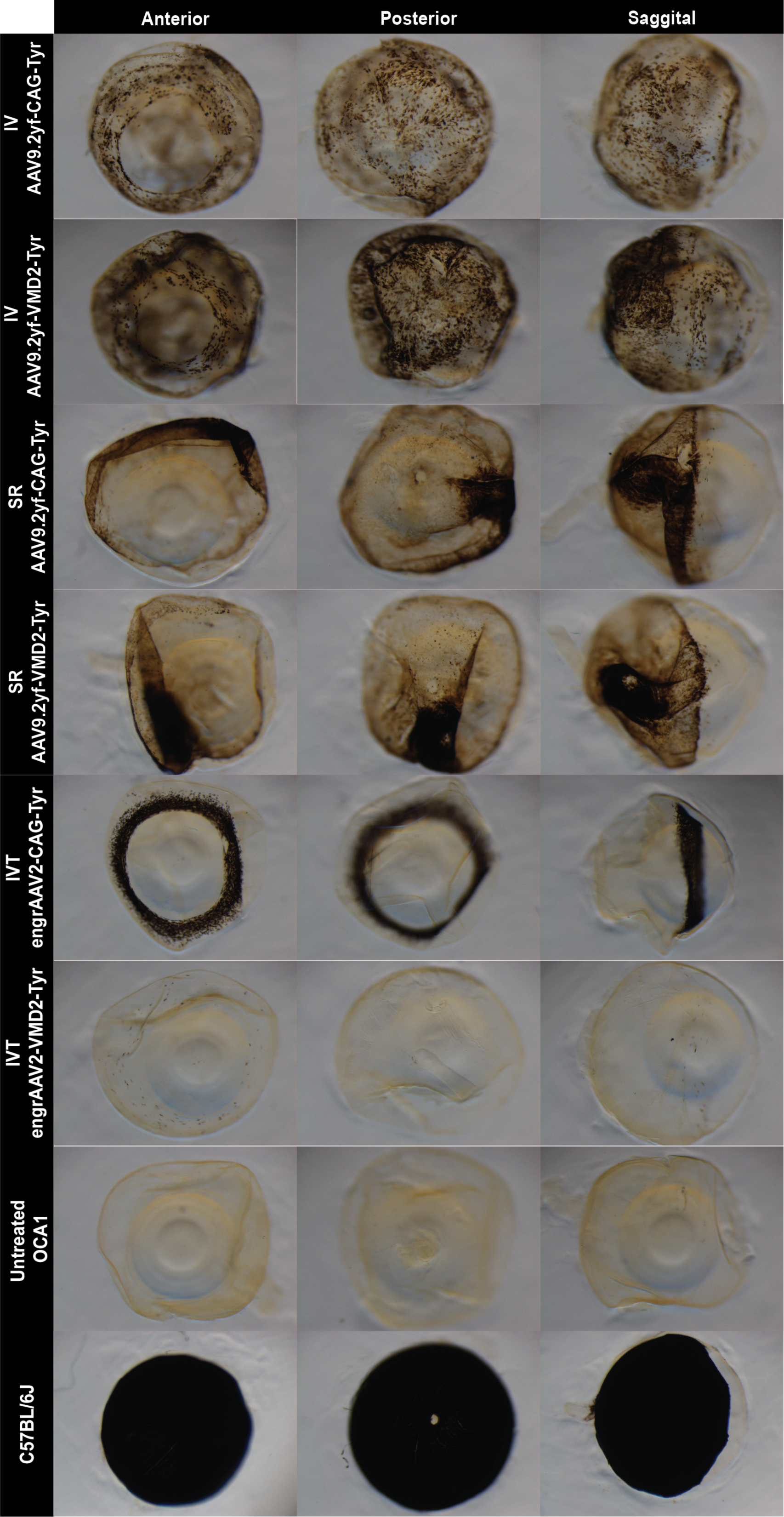
Whole eyes cleared using melanin-sparing tissue clearing techniques demonstrate that anterior and posterior segment pigment patterning is dependent on viral delivery method. Intravenous AAV9.2yf-*Tyr* resulted in both iris and widespread posterior segment pigmentation, with AAV9.2yf-VMD2-*Tyr* producing qualitatively more extensive posterior segment pigment than the ubiquitous vector. Subretinal AAV9.2yf-*Tyr* resulted in concentrated posterior pigment deposition localized to the injection site, with VMD2 and CAG constructs resulting in similar pigmentation levels. No iris pigmentation is observed following subretinal injection. Intravitreal injection with the ubiquitous construct resulted in dense iris pigmentation, while VMD2-*Tyr* resulted in sparse iris pigment deposition. Posterior segment pigmentation is negligible in intravitreally-treated eyes. Untreated OCA1 and C57Bl/6J controls serve as references for non-pigmented and fully pigmented eyes, respectively.

To assess the long-term stability of pigment expression following AAV-*Tyr* transduction, we performed serial *in vivo* imaging at one month and again greater than five months post-treatment. Pigmentation observed at one month was stably maintained over time, with a modest increase in posterior segment pigmentation coverage observed in aged mice, suggesting ongoing melanin production (Figure 4).

**Figure 4.**
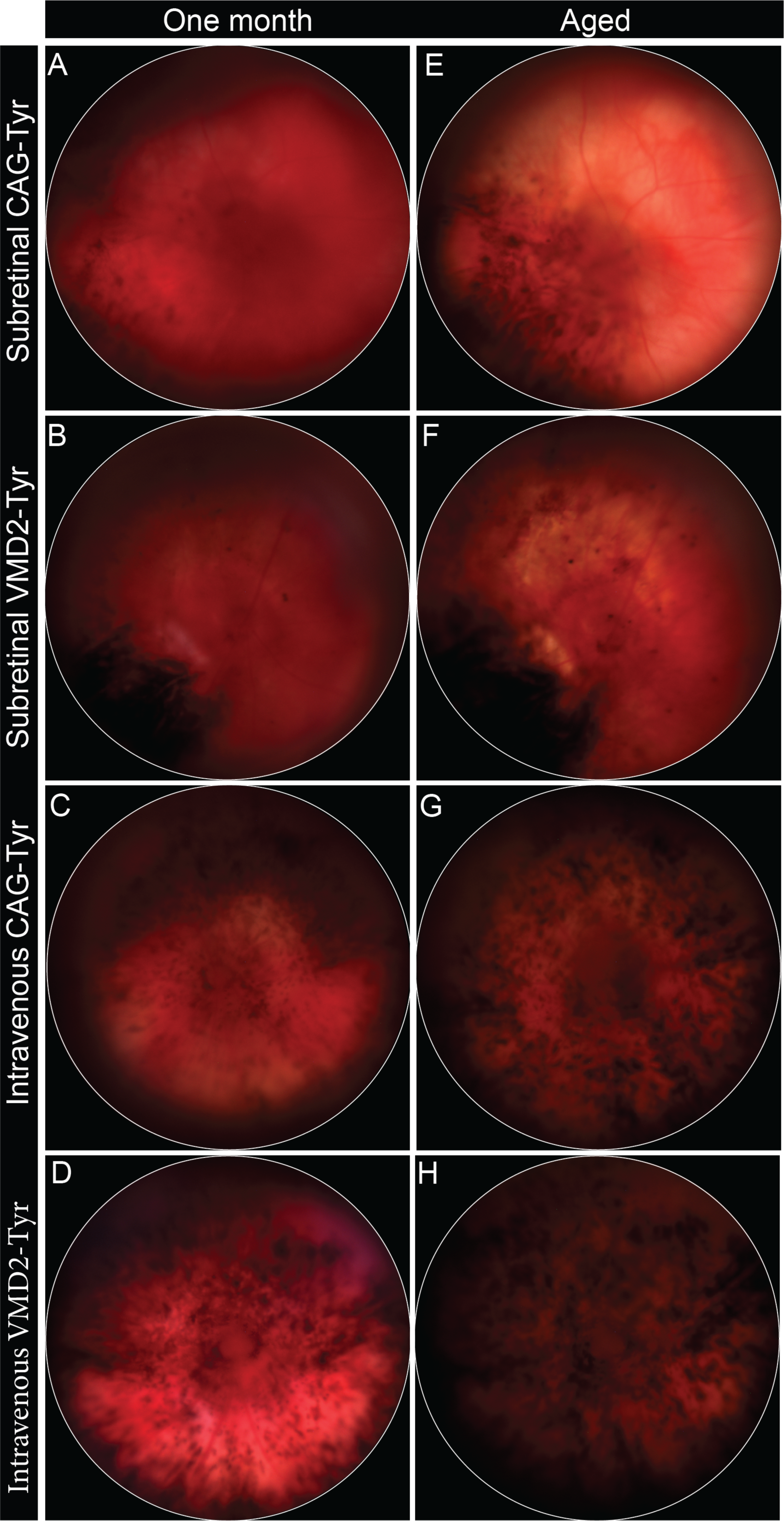
Posterior segment pigmentation increased with age in subretinally and intravenously treated OCA1 mice. (**A-D**) *In vivo* fundus images acquired one-month following subretinal (**A-B**) and intravenous (**C-D**) AAV9.2yf-*Tyr* delivery. (**E-H)** Serial fundus images obtained from the same mice aged 7-8 months (**E-F**) and 5-6 months (**G-H**) post viral delivery demonstrate increased posterior segment pigment deposition and area of distribution in both subretinal and intravenous treatment groups.

Both *in vivo* and whole cleared eye imaging demonstrated increased posterior segment pigmentation in VMD2-*Tyr* treated eyes qualitatively. To compare the efficacy of CAG-*Tyr* and VMD2-*Tyr* constructs head-to-head, we quantified the pigmented surface area in wholemounts containing RPE and choroid, which appear indistinguishable at the imaging resolution used (Figure 5A). Intravenous delivery yielded the highest pigmented surface area, with 56% ± 11 and 65% ± 1.7 of the RPE/choroid surface pigmented in AAV9.2yf-CAG-*Tyr* and AAV9.2yf-VMD2-*Tyr* treated eyes, respectively. Although VMD2-driven *Tyr* expression resulted in greater pigmentation on average, this difference did not reach statistical significance (p= 0.3217). The subretinal method of injection resulted in substantially less RPE/choroid area pigmented, averaging 29% ± 8.5 and 29% ± 12 in CAG-*Tyr* and VMD2-*Tyr* groups, respectively. Eyes treated via intravitreal injection displayed negligible posterior segment pigmentation for both constructs (Figure 5B).

**Figure 5.**
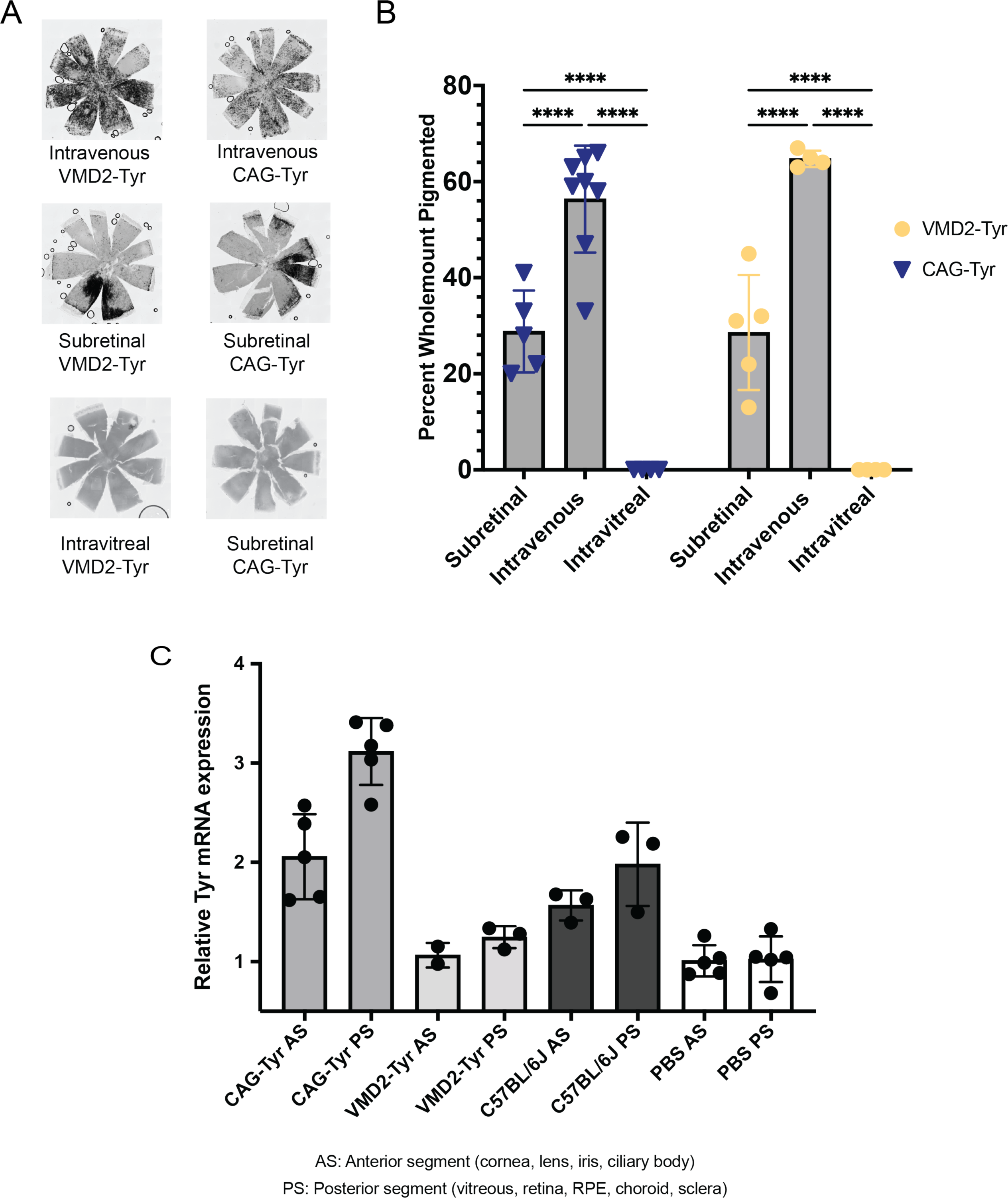
Quantitative analysis of ocular pigment restoration and *Tyr* expression. (**A**) Representative posterior segment wholemounts containing retina, RPE, and choroid tissue obtained from mice aged 5-6 months after intravenous injection, 7-8 months after subretinal injection, and 5-6 months after intravitreal AAV-*Tyr* injection. (**B**) Quantification of pigmented surface area in CAG-*Tyr* and VMD2-*Tyr* treatment groups across intravenous, subretinal, and intravitreal delivery methods. Data were analyzed by two-way ANOVA with Šídák’s multiple comparisons test. Non-significant comparisons not shown. ****p<0.0001 (**C**) Relative *Tyr* mRNA expression in anterior segment (cornea, lens, iris, ciliary body) and posterior segment (vitreous, retina, RPE, choroid, and sclera) tissues four weeks after P1 intravenous AAV9.2yf-CAG-*Tyr* or AAV9.2yf-VMD2-*Tyr* injection. The ddCT method was used to calculate fold difference in *Tyr* expression normalized to sham-treated OCA1 control. PBS=sham-treated OCA1 control. C57BL/6J=fully pigmented control. Data were analyzed by Kruskal-Wallis test with Dunn’s multiple comparisons test. Non-significant comparisons not shown. *p<0.05; ** p<0.01. All error bars represent standard deviation.

To investigate whether differences in *Tyr* transgene expression could explain the observed trend of increased posterior segment pigmentation following intravenous VMD2-*Tyr* versus CAG-*Tyr* delivery, we quantified relative Tyr mRNA transcript abundance in ocular tissue following P1 intravenous AAV9.2yf-*Tyr* injection (Figure 5C). AAV9.2yf-CAG-*Tyr* resulted in 2.06-fold and 3.17-fold increase in *Tyr* transcript levels in anterior and posterior segment tissues, respectively, compared to sham-treated OCA1 controls (p=0.0484; p=0.0017). Conversely, intravenous AAV9.2yf-VMD2-*Tyr* results in a minimal increase Tyr transcript abundance, with a 1.07-fold and 1.25-fold increase in *Tyr* transcript levels in anterior and posterior segment tissue, respectively, compared to sham OCA1 control (p>0.9999; p>0.9999). Together, these findings indicate that the VMD2 promoter drives lower overall transcript expression but achieves comparable or greater pigment restoration, potentially related to more targeted expression in pigment-producing tissues.

To determine whether visible pigmentation correlated with melanogenesis at the ultrastructural level in RPE and choroid tissues, we assessed melanosome maturation through transmission electron microscopy (TEM) eight-weeks post-intravenous delivery with AAV9.2yf-CAG-*Tyr* and AAV9.2yf-VMD2-*Tyr*. Melanosome biogenesis is divided into four stages based on ultrastructural features. Stage I and II represent early pre-melanosomes with lysosomal-like morphology and structural fibrils that serve as scaffolds for eventual melanin deposition. Stage III mature melanosomes are partially pigmented but retain visible fibrillar architecture. Stage IV melanosomes are fully pigmented and appear uniformly electron dense, with the underlying melanized fibrillar scaffold obscured from view.

High-magnification TEM of the RPE revealed a substantial reduction in the average number of melanosomes (stage I-IV) in untreated OCA1 mice compared to fully pigmented C57BL/6J controls (0.17 ± 0.071 vs. 0.73 ± 0.064 melanosomes/um^2^, respectively) (Figure 6). These findings are consistent with previous reports of impaired melanosome biogenesis in OCA1 mice^23,30,31^. Eyes treated with intravenous AAV9.2yf-CAG-*Tyr* and AAV9.2yf-VMD2-*Tyr* displayed a similar reduction in the average number of RPE melanosomes/um^2^ (0.31 ± 0.078 and 0.13 ± 0.046, respectively). However, AAV9.2yf-*Tyr* increased the mean number of mature RPE melanosomes, with 0.10 ± 0.051 and 0.037 ± 0.018 stage III/IV melanosomes/um^2^ observed in CAG and VMD2-*Tyr* groups, compared to untreated OCA1 mice which predictably displayed no stage III/IV melanosomes. Among the melanosomes present in the RPE, 34 ± 13%, 29 ± 7.5%, and 95 ± 1.5% were pigmented in CAG-*Tyr,* VMD2-*Tyr*, and C57BL/6J groups, respectively.

**Figure 6.**
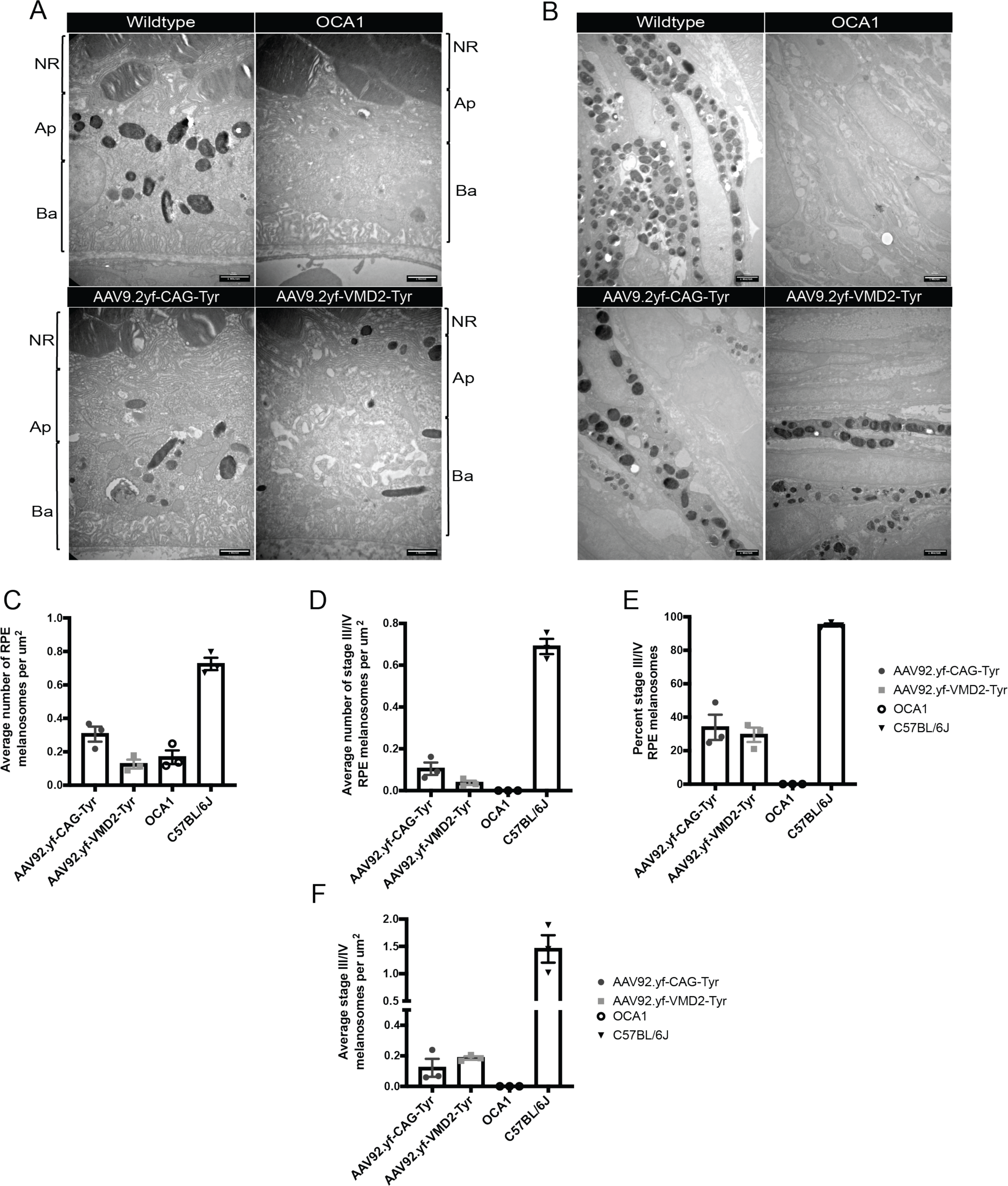
AAV9.2yf-CAG-Tyr and AAV9.2yf-VMD2-*Tyr* restored melanosome pigment deposition in RPE and choroid. (**A**) Transmission electron microscopy (TEM) images of RPE from AAV9.2yf-CAG-*Tyr* and AAV9.2yf-VMD2-*Tyr*-treated mice, collected 8-weeks after intravenous injection. Age-matched wildtype and OCA1 controls are shown for comparison. Images acquired at 20,000x magnification. NR = neural retina; Ap = apical RPE; Ba = basal RPE. (**B**) TEM images of choroid from the same treatment groups and time points. Images acquired at 15,000x magnification. (**C**) Quantification of total (stage I-IV) RPE melanosomes/um^2^, averaged across 18-20 TEM microphotographs per biological replicate. (**D**) Quantification of mature (stage III/IV) melanosomes/um^2^. (**E**) Percentage of mature (stage III/IV) melanosomes relative to the total RPE melanosome count. (**F**) Quantification of mature (stage III/IV) melanosomes/um^2^ in the choroid, averaged across 6 TEM microphotographs imaged at 2,000x magnification per biological replicate. Scale bars represent 1 um. All error bars represent standard deviation.

Choroid melanosome pigmentation was evaluated at lower magnification to assess a wider tissue area. This permitted quantification of stage III/IV melanosomes, but precluded distinction of pre-melanosomes from other lysosomal structures. AAV9.2yf-CAG-*Tyr* and AAV9.2yf-CAG-*Tyr* restored melanosome pigmentation in choroidal melanocytes, which was absent in untreated controls. CAG and VMD2-*Tyr* groups demonstrated 0.12 ± 0.10 and 0.19 ± 0.020 stage III/IV melanosomes/um^2^, respectively, compared to 1.5 ± 0.43 stage III/IV melanosomes/um^2^ in C57BL/6J choroid.

To assess potential off-target AAV-*Tyr* expression, we quantified relative *Tyr* transcript abundance in two non-ocular tissues: the liver, a common site for AAV transduction-related adverse events^32,33^, and brain subcortex, including the substantia nigra, a region that is vulnerable to deleterious effects following ectopic *Tyr* expression related to dopaminergic pathway involvement^34^. RT-qPCR analysis revealed a 255-fold and a 34-fold increase in hepatic *Tyr* transcript abundance in mice treated with intravenous AAV9.2yf-CAG-*Tyr* and AAV9.2yf-VMD2-*Tyr*, respectively, compared to untreated OCA1 controls (Figure S3). Intravenous AAV9.2yf-CAG-*Tyr* led to variable *Tyr* expression within brain subcortex, with an average 36-fold increase in relative transcript abundance (Figure S3). Conversely, intravenous AAV9.2yf-VMD2-*Tyr* resulted in a 2-fold increase in *Tyr* transcript expression in subcortex. Subretinal and intravitreal methods of injection resulted in negligible hepatic and subcortex *Tyr* expression.

Based on the superior pigment restoration achieved with intravenous delivery route and the reduced off-target expression conferred by the VMD2 promoter, we selected intravenous AAV9.2yf-VMD2-*Tyr* as our lead therapeutic candidate for functional validation. We quantified photophobic behavior using the light-dark box assay (Figure 7A). Previous studies demonstrated that OCA1 mice exhibit light avoidance at high luminosities, mirroring the photophobia experienced by OCA1 patients^35^. At low lighting levels (0 and 150 lux), intravenous AAV9.2yf-VMD2-*Tyr*-treated and untreated mice spent a similar percentage of time in the lit compartment. However, at high luminosity (1500 lux), AAV9.2yf-VMD2-*Tyr*-treated mice spent significantly more time in the lit compartment compared to untreated controls (45.0 ± 12.6% vs 22.0 ± 3.3%, p=0.0104) (Figure 7C). The number of dark-to-light compartment crossover events did not differ between groups at any luminosity, indicating exploratory drive was intact in both groups (Figure 7D).

**Figure 7.**
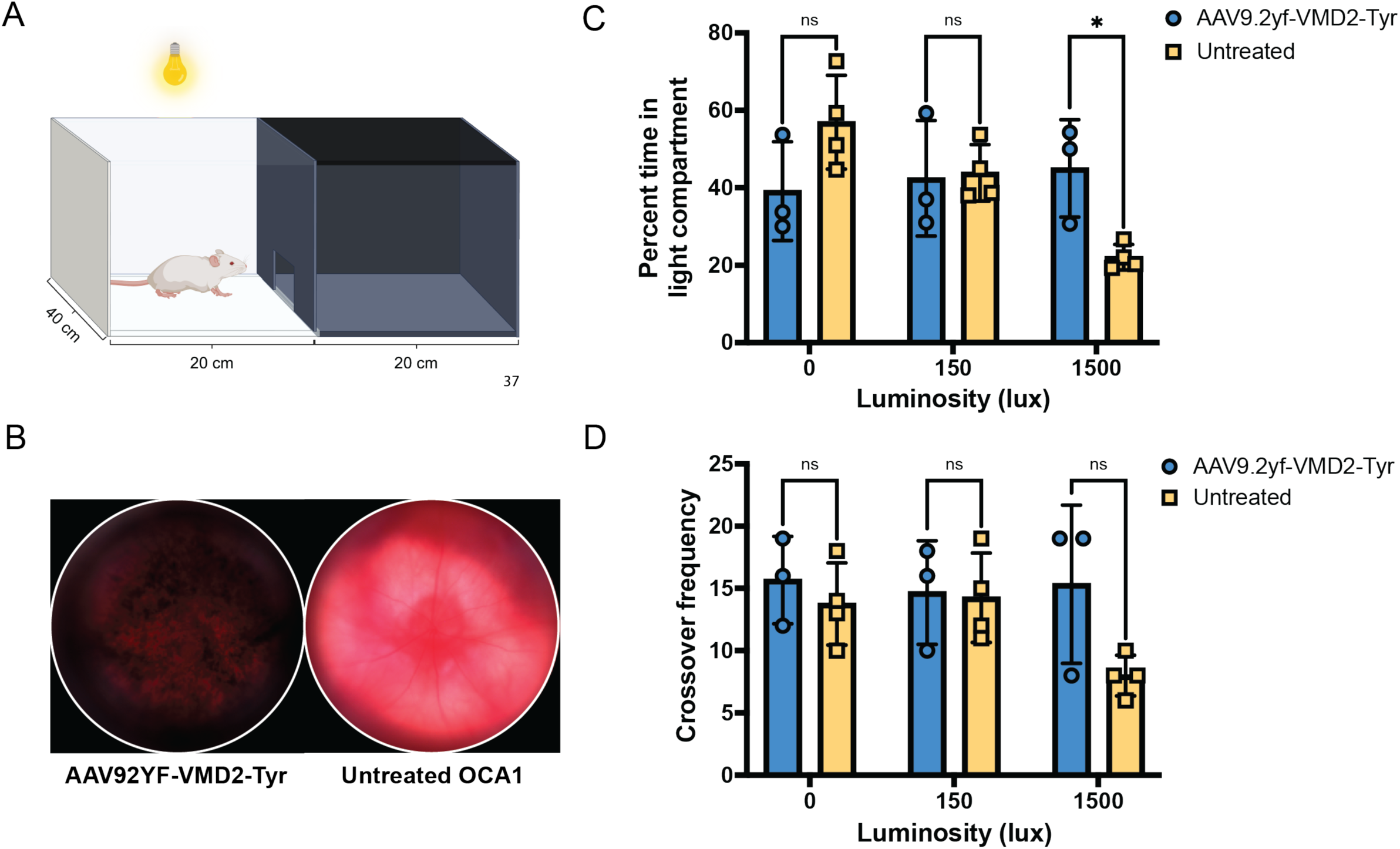
Intravenous AAV9.2yf-VMD2-*Tyr* improved light tolerance. (**A**) The dual chamber behavioral arena used to quantify photophobic behavior includes a dark, light-proof compartment (right) and illuminated compartment (left) that mice were able to freely pass between; compartment preference was tracked using a single overhead IR-sensitive camera at 0, 150, and 1500 lux (**B**) Representative fundus images depicting posterior segment pigment restoration in an intravenous AAV9.2yf-VMD2-*Tyr*-treated mouse compared to an untreated, sham (intravenous PBS) OCA1 control (**C**) Light aversion was quantified via the percent of each five minute trial spent in the light-exposed compartment at 0, 150, and 1500 lux (**D**) Crossover frequency between chambers was measured at 0, 150, and 1500 lux. Data were analyzed by 2-way ANOVA with Šídák’s multiple comparisons test. *p<0.05. All error bars represent standard deviation.

## DISCUSSION

The success of an AAV-mediated gene replacement therapy depends on defining key delivery parameters, including the method of viral administration and the cell specificity of transgene expression. In the present study, we evaluated three AAV-*Tyr* delivery methods, each chosen for their hypothesized ability to maximize viral particle transduction in pigmented ocular tissue, in combination with two *Tyr*-expressing constructs with varying RPE-specificity, to determine the optimal *Tyr* replacement strategy for ocular pigment restoration. This study represents the first systematic comparison of viral delivery route and promoter specificity for effective ocular pigmentation and functional vision rescue in an OCA1 model.

*In vivo* fundus imaging, whole eye imaging, and posterior segment wholemount pigment analysis demonstrated that intravenous AAV9.2yf-*Tyr* delivery resulted in the most widespread pigmentation restoration in both anterior and posterior segments. The pigmentation pattern observed following intravenous AAV9.2yf-*Tyr* injection likely reflects the AAV9.2yf capsid’s ability to traverse immature retinal vascular endothelium via transcytosis^27^, enabling homogenous RPE and uveal melanocyte transduction. In contrast, subretinal AAV9.2yf-*Tyr* delivery resulted localized in pigment restoration surrounding the injection site, consistent with previous studies demonstrating limited lateral AAV spread beyond the boundary of subretinal neurosensory detachment^36^. The intravitreal method of vector delivery was unsuccessful in targeting RPE and choroid at the tested dose (3.8e9 vg). However, robust iris pigmentation was observed in intravitreal CAG-*Tyr* treated eyes, likely due to anterior viral drainage which facilitated iris melanocyte transduction. Taken together, these data support the conclusion that early, intravenous AAV9.2yf-*Tyr* administration results in the most effective ocular pigment restoration in this OCA1 model.

In addition to identifying the optimal viral delivery method, selecting a promoter that results in efficient, on-target transgene expression in cell type(s) of interest is critical for efficacy and safety. We evaluated pigment restoration using a ubiquitous construct (CAG-*Tyr*) compared to an RPE-specific construct (VMD2-*Tyr*), driven by MITF-binding elements that are primarily expressed in RPE^28^. Whole cleared eyes and TEM reveal iris and choroid pigmentation, respectively, in addition to RPE, suggesting the truncated VMD2 promoter used in this study is active within cranial neural crest-derived melanocytes as well as RPE. Surprisingly, intravenous AAV9.2yf-VMD2-*Tyr* but not intravitreal engrAAV2-VMD2-*Tyr* resulted in significant iris pigment rescue. This phenomena may be related to capsid-promoter interactions which alter cell specificity, where variable transduction profiles can be observed in AAV9 versus AAV2-based capsids despite the use of a consistent promoter/transgene^37^.

Quantitative wholemount analysis revealed VMD2-*Tyr* resulted in similar posterior segment pigmentation to CAG-*Tyr* construct across all delivery methods. These findings are in contrast to a recently published study, which found VMD2-*TYR* to result in minimal to no pigment production when delivered via an AAV8 vector subretinally to OCA1 mice^38^. Our results, which demonstrate VMD2-*Tyr* results in pigment restoration that is comparable to a ubiquitous promoter suggests the AAV9.2yf-VMD2-*Tyr* vector used herein results in superior RPE transduction as compared to AAV8.

Despite comparable pigmentation rescue between CAG-*Tyr* and VMD2-*Tyr* constructs, intravenous AAV9.2yf-CAG-*Tyr* resulted in higher *Tyr* mRNA expression in anterior (cornea, iris, ciliary body) and posterior segment (retina, RPE, choroid, sclera) tissue compared to the sham OCA1 group. Further, intravenous AAV9.2yf-CAG-*Tyr* led to notably greater *Tyr* expression in liver and subcortex tissue of all AAV-*Tyr* approaches tested, highlighting unintended off-target expression. Hepatic overexpression of AAV transgene has been associated with hepatotoxicity^33^ and ectopic *TYR* expression in substantia nigra has been linked to melanin-mediated dopamine signaling disruption resulting in Parkinsonian symptoms in rodents^34^. Together, our findings underscore the importance of promoter choice and support VMD2-*Tyr* as a safer alternative to ubiquitous expression constructs for ocular pigment restoration in OCA1.

Ultrastructural analysis revealed partial melanogenesis rescue in RPE and choroid following intravenous AAV9.2yf-CAG-*Tyr* and AAV9.2yf-VMD2-*Tyr* delivery. RPE melanosome biogenesis begins during embryogenesis, at approximately E10.5 in mice, and is largely complete by postnatal developmental stages^39^. OCA1 RPE fail to form mature melanosomes and demonstrate an overall reduction in the number of melanosomes in both developing and developed RPE^23,30,31^. Our data adds to the body of literature documenting reduced melanosome density in OCA1 RPE and demonstrate melanosome pigment deposition and maturation may be achieved using intravenous AAV9.2yf-*Tyr*. Post-natal pigmentation rescue, as was performed in this study, may be limited by inborn melanosome production defects resulting in reduced melanosome availability in developed OCA1 RPE.

While restoration of OCA-related developmental anomalies are not likely using a gene therapy administered postnatally, after retinal development is complete, we postulate that AAV-mediated *TYR* replacement therapy will improve pigment-related vision. We hypothesize that improvement in visual function will correlate with ocular pigment restoration in patients with OCA1, mediated by reduced light scatter, glare, and photophobia. Here, we have demonstrated that OCA1 mice treated with intravenous AAV9.2yf-VMD2-*Tyr* spend more time occupying the lit compartment of a light-dark chamber at a high luminosity (1500 lux) compared to untreated controls. Light compartment avoidance at high but not ambient lighting conditions was established as a measure of photophobic behavior in OCA1 mice^35^. OCA1 mice treated intravenously with our leading vector candidate, AAV9.2yf-VMD2-*Tyr*, demonstrated reduced photophobic behavior while untreated controls display expected high-intensity light avoidance. Together, these results indicate that AAV-mediated pigment restoration mitigates a debilitating functional feature of OCA1 and may confer visual benefit for OCA1 patients.

Our data indicate that early, intravenous AAV9.2yf-*Tyr* delivery results in the greatest posterior segment pigmentation in OCA1 mice. Key to the success of the intravenous approach tested herein bypassing the blood-retina barrier, which remains permissive to AAV9.2yf transcytosis until approximately P7^27^. Translating these findings to the clinic will require innovative strategies to bypass the human blood-retina barrier and achieve efficient gene delivery to the RPE. In humans, the analogous stage of vascular development occurs between gestational weeks 14 and 18^40^. A systemic, *in utero* approach may be considered in the future to achieve both pigmentation and retinal development rescue in the OCA1 disease context. Alternatively, future studies are warranted to investigate the efficacy of an alternative AAV-*Tyr* approach, such as the suprachoroidal injection technique. Suprachoroidal viral vector delivery is noninvasive and deposits AAV in the highly vascular suprachoroidal space, resulting in widespread RPE transduction in non-human primate eyes^41^. The suprachoroidal approach is currently used in clinical practice and represents a translational alternative to systemic intravascular AAV-*Tyr* delivery.

In this study, we defined critical parameters for the development of an AAV-based gene therapy for OCA1. We demonstrate that delivery route and transgene expression specificity are key determinants of therapeutic efficacy and safety. Collectively, our findings identify intravenous, RPE-specific AAV-*Tyr* as the leading gene therapy strategy for achieving widespread ocular pigmentation and mitigating photophobic behavior. By restoring ocular pigmentation and reducing photosensitivity in OCA1 mice, we establish a meaningful therapeutic benefit to postnatal TYR replacement. These results lay the foundation for future clinical investigation of AAV-*TYR* therapies aimed at alleviating painful photosensitivity and improving visual outcomes in patients with OCA1.

## MATERIALS AND METHODS

### Animals

All procedures were performed in accordance with the Association for Research in Vision and Ophthalmology Statement for the Use of Animals in Ophthalmic and Vision Research and approved by the Institutional Animal Care and Use Committee at the University of Pittsburgh. B6(Cg)-Tyr^c-2J^/J (OCA1) and C57BL/6J mice were obtained from the Jackson Laboratory and bred at the University of Pittsburgh. Animals were housed in a standard 12/12-hr light/dark cycle and were provided food, water, and enrichment. Male and female mice were used for all experiments.

### AAV Packaging and Purification

AAV vectors were produced in 293AAV cells (Cell Biolabs, San Diego, CA, USA) using the previously described triple transfection method^42^. Prior to packing, all plasmids were sequence-verified through whole plasmid sequencing (Plasmidsaurus, Louisville, KY, USA) and ITR integrity was assessed via restriction digest. Virus was purified from crude cell lysate through serotype-specific affinity chromatography^43,44^, buffer exchanged, and concentrated using Amicon Ultra-15 Centrifugal Filter Units in D-PBS. Purified AAVs were titered using a quantitative PCR method with ITR-binding primers (abm, Richmond, BC, Canada). Vectors were normalized to a concentration of 2.5e12 vg/mL (± 1.5e12 vg/mL).

### AAV Delivery

#### Intravenous AAV injection in P1 pups

Pups were removed from their home cage and placed on a 37C warming pad along with bedding. Pups were placed on ice for anesthesia/analgesia for one minute, or until loss of response to tactile stimuli was observed. The facial vein was identified in anesthetized pups using a dissecting microscope. 50 uL of virus at a concentration of 2.5e12 vg/mL or PBS (control) was injected via the facial vein using a 31G, 8mm insulin syringe (BD). Injection success was confirmed by observed clearing of the facial vein and branching vessels immediately after intravenous flush. Pups were recovered using a warming pad and returned to their cages in stable condition.

#### Subretinal AAV injection in 1-month-old mice

Mice were anesthetized using ketamine (90 mg/kg) and xylazine (10 mg/kg) delivered via intraperitoneal injection. Anesthetized mice were placed on a 37C warming pad. Mydriasis was achieved bilaterally using 1% Tropicamide and 2.5% Phenylephrine, 0.5% Proparacaine was used for analgesia, and corneal moisture was maintained using Genteal ophthalmic ointment. A 12 mm glass coverslip (Fisher Scientific) was placed on the surface of the eye and used to visualize the retina with the assistance of a dissecting microscope. A sclerotomy was performed 0.5 mm posterior to the limbus using a 31G hypodermic needle (BD) to achieve posterior segment access. A 33G blunt needle attached to a10 uL Hamilton syringe was passed through the sclerotomy site and used to perform a retinotomy and subretinal injection ∼4 disc diameters temporal to the optic nerve head. 1.5 uL of AAV at a concentration of 2.5e12 vg/mL (right eye) or PBS (left eye) was injected subretinally in each mouse. Mice were recovered on a heating pad and returned to their cage after fully awakening.

#### Intravitreal AAV injection in 1-month-old mice

Mouse anesthesia, mydriasis, analgesia, and sclerotomy were performed as described above. A 33G blunt needle attached to a 10 uL Hamilton syringe was passed through the sclerotomy site, positioned above the optic nerve head, and used to deliver 1.5 uL of AAV at a concentration of 2.5e12 vg/mL (right eye) or PBS (left eye) into the vitreous cavity of each mouse. Mice were recovered on a heating pad and returned to their cage after fully awakening.

### Fundus Photography

Prior to imaging, mice were anesthetized using ketamine (90 mg/kg) and xylazine (10 mg/kg) delivered via intraperitoneal injection. Mydriasis, ocular analgesia, and corneal moisture were achieved using the methods described above. Iris and retinal images were obtained using the Pheonix Micron IV fundus camera system for rodents. Mice were recovered on a heating pad and returned to their cage after fully awakening.

### Whole Eye Clearing

Eumelanin-sparing tissue clearing techniques were used to visualize anterior and posterior segment pigmentation^29^. Immediately after euthanasia, eyes were enucleated and placed in 4% PFA. Eyes were fixed at room temperature for one hour followed by 4C fixation overnight. Fixed eyes were successively dehydrated in 20%, 40%, 60%, 80% and 100% methanol and delipidated in 66% dichloromethane in methanol overnight. Delipidated tissues were cleared in BABB (1:1 benzyl benzoate and benzyl alcohol) for at least 24 hours prior to imaging.

### Wholemount Tissue Preparation and Pigment Quantification

Immediately after euthanasia, eyes were enucleated and placed in Hartmann’s fixative. Eyes were fixed at room temperature for one hour followed by 4C fixation overnight. Eyes were transferred to 70% ethanol in deionized water for dissection. The anterior segment was removed and eight full-thickness relief cuts were made in the retina, RPE, choroid, and sclera to encourage the tissue to lay flat. Flattened tissue was mounted on a slide using gelvatol hardening medium. Brightfield images were acquired on the Evident Slideview VS200 using a 0.4NA 10x UPlanXApo objective and Evident VS200 ASW 3.4.1 software. Automatic z-range was used to create an Extended Focus Image (EFI) with 2.36um steps. Pigment was quantified in 8-bit images (intensity range 0-255) as the tissue pixel area with intensity values less than 90% of the median background intensity, expressed as a percentage of total tissue area. Visualizations were generated in Python [3.12.3] using Jupyter Notebooks executed within Visual Studio Code [v1.102.3] and Napari.

### Transmission Electron Microscopy

Immediately after euthanasia, eyes were enucleated and placed in 4% paraformaldehyde/2.5% glutaraldehyde for overnight fixation. Samples were processed with osmium tetroxide, stained with Uranyless and lead aspartate (2 minutes each), embedded, and sectioned as previously described^45^. TEM images were acquired using a JEOL 1400-PLUS TEM transmission electron microscope at 80 kV. Melanosome developmental stages were identified using established ultrastructural criteria^46^. Stage I melanosomes were identified as membrane-bound organelles containing vesicular and irregular fibrillar substructures. Stage II melanosomes resembled stage I but contained organized fibrillar striations in concentric arrays. Fibrillar melanin deposition was evident in stage III melanosomes, which were distinguishable from uniformly electron dense stage IV melanosomes. Melanosomes were counted manually and expressed relative to tissue area (um^2^). Image analysis was performed using ImageJ.

### Quantitative real time-polymerase chain reaction (qRT-PCR)

Whole RNA was extracted from eye, liver, and brain tissues using a FastPrep-24 Homogenizer (MP Biomedicals) and the RNEasy Mini Kit (Qiagen). Samples were DNase-treated using a TURBO DNA-free kit (Thermo Fisher) and purified using the Monarch Spin RNA Cleanup Kit (NEB). RNA quality was assessed through spectrophotometry and Qubit RNA quantification. cDNA synthesis was performed using the Verso cDNA Synthesis Kit (Thermo Scientific) using a 3:1 blend of random hexamers and anchored olig-dT primers. qPCR was performed using KAPA SYBR® FAST (Roche), Tyr primers (FWD: 5’ AGC CTG TGC CTC CTC TAA; REV: 5’ AGG AAC CTC TGC CTG AAA), and GAPDH primers (FWD: 5’ CAT CAC TGC CAC CCA GAA GAC TG; REV: 5’ ATG CCA GTG AGC TTC CCG TTC AG).

### Immunohistochemistry

Immediately after euthanasia, eyes were enucleated and placed in 4% paraformaldehyde for one hour and room temperature followed by overnight fixation at 4C. Eyes were transferred to PSB prior to cryoprotection in 15% and 30% sucrose/PBS solutions at RT for at least 30 minutes each. Tissue was transferred to 1:1 30% sucrose:OCT compound (Fisher Scientific) prior to embedding in 100% OCT compound.12 uM frozen sections containing optic nerve head were collected on poly-lysine coated glass slides and stored at -80C until immunolabeling was performed. Cryosections were rehydrated with PBS for 10 minutes, blocked with 10% normal goat serum, 1% BSA, and 0.5% Triton X-100 in PBS for one hour. Tyrosinase Polyclonal Antibody (Bioss) was diluted in 1:200 in blocking buffer and incubated for 12-16 hours at 4C. Washed tissue was incubated with donkey anti-rabbit Alexa IgG (H+L) Highly Cross-Adsorbed Secondary Antibody, Alexa Fluor™ 647 (Invitrogen) for 1 hour at 4C in the dark. Hoescht 33342 was used to label nuclei and was diluted 1:2000 in PBS and incubated for 30 minutes at room temperature in the dark. Slides were mounted with Vectashield mounting medium with DAPI (Thermo Fisher) and imaged using an Olympus Fluoview 1000 confocal microscope.

### Light-Dark Photophobia Assay

A light-dark assay was conducted as previously described^35,47–49^ using an dual-chamber arena containing a 40 × 20 × 30 cm light-protected compartment and a 40 × 20 × 30 cm compartment exposed to overhead lighting. Subcompartments were divided by a light-protected insert that contained a 7.5 × 3.75 cm opening to permit locomotion between compartments. Light compartment luminosity was confirmed using a luxometer (Med Associates). The mice used in this assay were previously acclimated to human handling and behavioral testing to reduce confounding anxiety. Mice were acclimated to the dual-chamber arena in 150 lux lighting conditions for two consecutive days, five minutes each day. Immediately following arena acclimation, photophobic behavior was assessed at 0, 150, and 1500 lux over a span of three days with one lighting condition tested per day. Prior to testing, mice were acclimated to the test lighting conditions for 45 minutes in in their home cage. Mice were placed individually in the testing arena and their behavior was recorded using a Basler acA1300-60gmNIR camera (Basler AG, Ahrensburg Germany) and analyzed with Noldus Ethovision XT 17.5 (Noldus Information Technology, Leesburg VA, USA) on a Lenovo ThinkPad (Lenovo, Bejing China) using Windows 10 (Microsoft, Seattle WA, USA).

### Statistical Analysis

Two-way Analysis of Variance (ANOVA) with Šídák’s multiple comparisons test was used to analyze wholemount pigmentation, taking into account method of delivery and viral construct variables. Relative *Tyr* mRNA expression in anterior and posterior segment was analyzed using Kruskal-Wallis test with Dunn’s multiple comparisons test. Two-way Analysis of Variance (ANOVA) with Šídák’s multiple comparisons test was used to analyze light-dark behavioral assay data, taking into account the effect of treatment and luminosity variables. All analyses were performed in GraphPad Prism 10 (Graphpad Software, La Jolla, CA, USA). All data were presented as means ± standard deviation. The criteria for significance were: p > 0.05, not significant; p < 0.05, *; p < 0.01, **; p < 0.001, ***; p < 0.0001, ****.

## DATA AVAILABILITY STATEMENT

Data generated in the current study are available from the corresponding author on reasonable request.

## ACKNOWLEDGEMENTS

Wholemount Slideview imaging was performed at the Center for Biologic Imaging at the University of Pittsburgh. We thank Mike Calderon for image acquisition and Dr. Katarzyna Kedziora for developing Python code for automated data analysis. All TEM images were acquired at the Center for Biologic Imaging at the University of Pittsburgh in collaboration with Dr. Simon Watkins. We thank Jonathan Franks for TEM sample preparation. Biorender was used for graphic generation. This work was supported by the NEI/NIH (F30EY036249 to ALP) and the National Organization for Albinism and Hypopigmentation (LB). This work was supported in part from an NIH CORE Grant (P30EY08098) and an unrestricted grant from Research to Prevent Blindness.

## AUTHOR CONTRIBUTIONS

Conceptualization, A.M.L.P and L.C.B; L.B.J performed TEM experiments and analyzed the data, W.G.K performed behavioral experiments, K.J.S performed IHC, A.M.L.P performed all other experiments; A.M.L.P analyzed the data and and wrote the manuscript; L.C.B and J.A.S provided resources; L.C.B, J.A.S, and A.M.L.P reviewed and revised the manuscript; All authors read and approved the final manuscript.

