## Supplemental Information for "Systemic, RPE-directed AAV-*Tyrosinase* therapy restores ocular pigmentation in an OCA1 mouse model"

### CAG

5’ cgttacataacttacggtaaatggcccgcctggctgaccgcccaacgacccccgcccattgacgtcaataatgacgta
tgttcccatagtaacgccaatagggactttccattgacgtcaatgggtggagtatttacggtaaactgcccacttggcagtac
atcaagtgtatcatatgccaagtacgccccctattgacgtcaatgacggtaaatggcccgcctggcattatgcccagtacat
gaccttatgggactttcctacttggcagtacatctactcgaggccacgttctgcttcactctccccatctcccccccctccccacc
cccaattttgtatttatttattttttaattattttgtgcagcgatgggggcgggggggggggggggcgcgcgccaggcggggcgg
ggcggggcgaggggcggggcggggcgaggcggagaggtgcggcggcagccaatcagagcggcgcgctccgaaa
gtttccttttatggcgaggcggcggcggcggcggccctataaaaagcgaagcgcgcggcgggcgggagcgggatcagc
caccgcggtggcggcctagagtcgacgaggaactgaaaaaccagaaagttaactggtaagtttagtctttttgtcttttatttc
aggtcccggatccggtggtggtgcaaatcaaagaactgctcctcagtggatgttgcctttacttctaggcctgtacggaagtg
ttacttctgctctaaaagctgcggaattgtacccgcggccgatcc 3’

### VMD2

5’ gaattctgtcattttactagggtgatgaaattcccaagcaacaccatccttttcagataagggcactgaggctgagagagga
gctgaaacctacccggcgtcaccacacacaggtggcaaggctgggaccagaaaccaggactgttgactgcagcccggtat
tcattctttccatagcccacagggctgtcaaagaccccagggcctagtcagaggctcctccttcctggagagttcctggcacag
aagttgaagctcagcacagccccctaacccccaactctctctgcaaggcctcaggggtcagaacactggtggagcagatcc
tttagcctctggattttagggccatggtagagggggtgttgccctaaattccagccctggtctcagcccaacaccctccaagaag
aaattagaggggccatggccaggctgtgctagccgttgcttctgagcagattacaagaagggaccaagacaaggactccttt
gtggaggtcctggcttagggagtcaagtgacggcggctcagcactcacgtgggcagtgccagcctctaagagtgggcaggg
gcactggccacagagtcccagggagtcccaccagcctagtcgccagacc 3’

**Figure S1 CAG and VMD2 promoter sequences.** Complete sequences for CAG and VMD2 promoters in the 5’ to 3’ orientation.

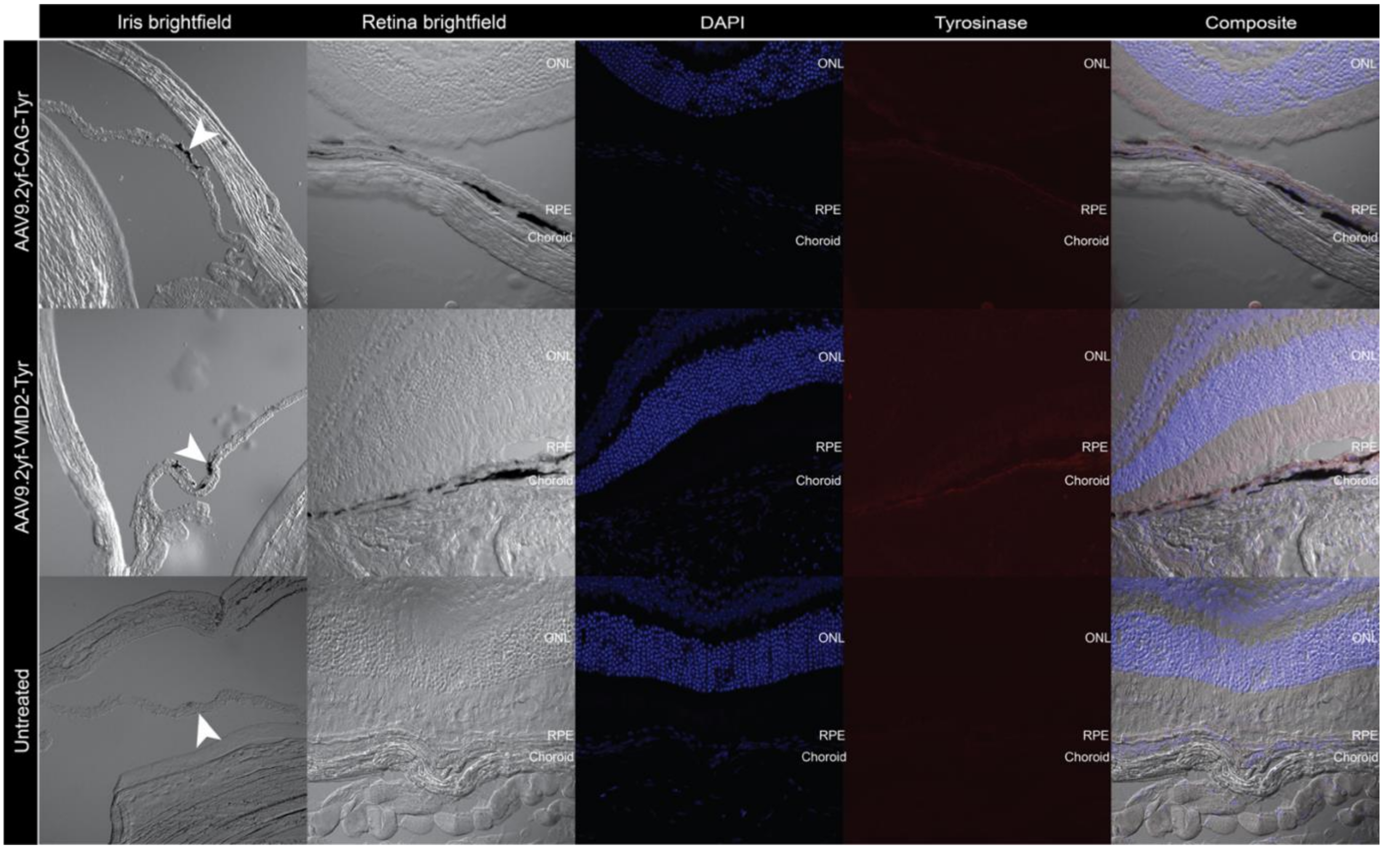


**Figure S2 Intravenous AAV9.2yf-*Tyr* restored Tyr expression and pigmentation in iris, RPE, and choroid.** Pigmentation was evident in iris (arrowhead), RPE, and choroid of animals treated with intravenous AAV9.2yf-CAG-*Tyr* and AAV9.2yf-VMD2-*Ty* but absent in untreated OCA1 tissue. Tyrosinase expression (red) correlates with pigmentation. Iris images were acquired at 20X magnification; RPE and choroid images were acquired at 40X magnification.

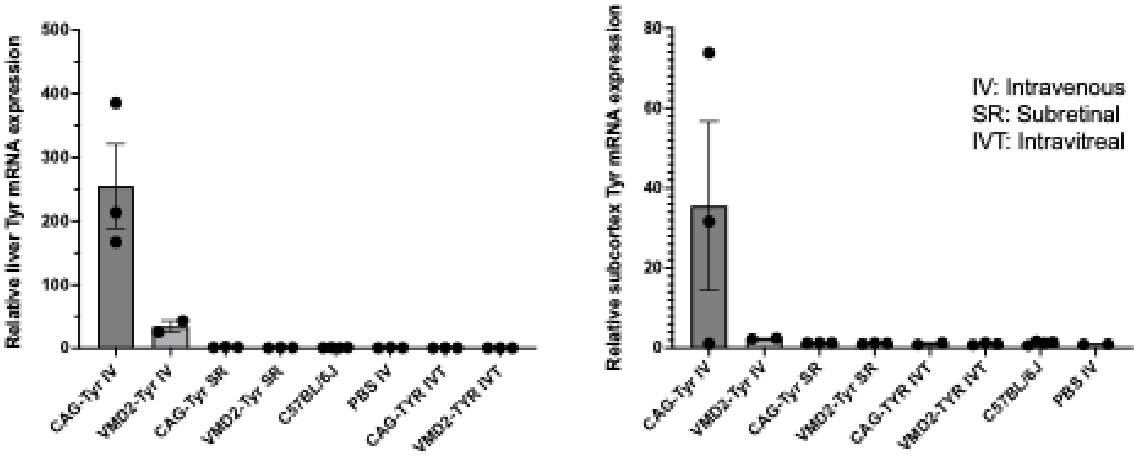


**Figure S3 Intravenous AAV9.2yf-CAG-*Tyr* resulted in off-target *Tyr* expression in liver and subcortex.** (A) Relative *Tyr* mRNA expression in liver following intravenous (IV), subretinal (SR), and intravitreal (IVT) delivery of rAAV-CAG-*Tyr* or rAAV-VMD2-*Tyr* (n=2-4). (**B**) Relative *Tyr* mRNA expression in brain subcortex, including substantia nigra, in the same treatment and injection groups. The ddCT method was used to calculate fold change in *Tyr* expression, normalized to sham-treated OCA1 control mice. PBS = P1 intravenous sham-treated OCA1 control; C57BL/6J = fully pigmented control. Each data point represents an individual biological replicate. RT-qPCR performed in duplicate. Error bars represent standard deviation of dCT values.

ß
